# Molecular determinants of SR-B1-dependent *Plasmodium* sporozoite entry into hepatocytic cells

**DOI:** 10.1101/2020.03.16.994731

**Authors:** Anne-Claire Langlois, Giulia Manzoni, Laetitia Vincensini, Romain Coppée, Carine Marinach, Maryse Guérin, Thierry Huby, Véronique Carrière, François-Loïc Cosset, Marlène Dreux, Eric Rubinstein, Olivier Silvie

## Abstract

Sporozoite forms of the malaria parasite *Plasmodium* are transmitted by mosquitoes and first infect the liver for an initial round of replication before parasite proliferation in the blood. The molecular mechanisms involved during sporozoite invasion of hepatocytes remain poorly understood. Two receptors of the Hepatitis C virus (HCV), the tetraspanin CD81 and the scavenger receptor class B type 1 (SR-B1), play an important role during the entry of *Plasmodium* sporozoites into hepatocytic cells. In contrast to HCV entry, which requires both CD81 and SR-B1 together with additional host factors, CD81 and SR-B1 operate independently during malaria liver infection. Sporozoites from human-infecting *P. falciparum* and *P. vivax* rely respectively on CD81 or SR-B1. Rodent-infecting *P. berghei* can use SR-B1 to infect host cells as an alternative pathway to CD81, providing a tractable model to investigate the role of SR-B1 during *Plasmodium* liver infection. Here we show that mouse SR-B1 is less functional as compared to human SR-B1 during *P. berghei* infection. We took advantage of this functional difference to investigate the structural determinants of SR-B1 required for infection. Using a structure-guided strategy and chimeric mouse/human SR-B1 constructs, we could map the functional region of human SR-B1 within apical loops, suggesting that this region of the protein may play a crucial role for interaction of sporozoite ligands with host cells and thus the very first step of *Plasmodium* infection.

**IMPORTANCE:** Malaria is caused by *Plasmodium* parasites and remains one of the deadliest parasitic diseases worldwide. The parasite is transmitted by a blood feeding mosquito and first invades the liver for an initial, obligatory and silent round of replication. The liver infection is an attractive target for antimalarial vaccine strategies, however the molecular mechanisms of parasite invasion of hepatocytes remain to be fully elucidated. Two hepatocyte surface proteins are known to be important for parasite entry into hepatocytes, the tetraspanin CD81 and the scavenger receptor class B type 1 (SR-B1). These receptors constitute independent gateways depending on the *Plasmodium* species. Here, we identified the structural determinants of SR-B1, an important hepatocyte entry factor for human-infecting *P. vivax*. This study paves the way toward a better characterization of the molecular interactions underlying the crucial early stages of infection, a pre-requisite for the development of novel malaria vaccine strategies.

## INTRODUCTION

Despite progress in malaria control over the last two decades, *Plasmodium* parasites continue to cause more than 200 million cases every year (1). After their inoculation into the skin by infected *Anopheles* mosquitoes, *Plasmodium* sporozoites rapidly migrate to the liver using gliding motility and cell traversal activity. Once in the liver, they first traverse hepatocytes before invading them and developing into exo-erythrocytic forms (EEFs), surrounded by a parasitophorous vacuole (PV) membrane. Inside the PV, they differentiate into thousands of merozoites, which are released into the blood circulation and invade red blood cells, provoking the symptomatic phase of the disease.

Several host and parasite factors implicated in sporozoite invasion have been identified but the underlying molecular interactions remain unknown. Human and murine parasites share similar invasion routes, with two distinct invasion pathways that depend on the tetraspanin CD81 or the scavenger receptor class B type 1 (SR-B1) (2–5). The human parasite *P. falciparum* and the murine parasite *P. yoelii* both require CD81 (3), whereas *P. vivax* enters human hepatocytes using SR-B1 (4). Interestingly, the murine parasite *P. berghei* can invade cells using either CD81 or, alternatively, a SR-B1-dependent route in the absence of CD81 (4). Whilst SR-B1 is the only known hepatocyte entry factor for *P. vivax* sporozoites, studying this parasite remains difficult, notably due to the limited access to infected mosquitoes. In this context, *P. berghei* provides an attractive model to investigate the role of SR-B1 during sporozoite infection.

SR-B1 is a highly glycosylated transmembrane protein that belongs to the CD36 family, which also includes CD36 and the lysosomal integral membrane protein 2 (LIMP-2). A tertiary structure of SR-B1 was predicted using LIMP-2 crystal structure as a template (6). SR-B1 possesses two transmembrane regions, cytoplasmic N- and C-termini, and a large extracellular domain constituted by a ß-strand tunnel topped by a helical bundle (6, 7). SR-B1 apical helices are involved in the binding of high density lipoproteins (HDLs) (8). The hydrophobic cavity traversing the entire protein is implicated in a selective lipid transfer with cholesteryl ester bidirectional exchanges between HDLs and the cell membrane (8, 9).

In this study, we show that murine SR-B1 poorly supports *P. berghei* infection as compared to its human counterpart. We took advantage of this functional difference to study the structural determinants of the SR-B1 receptor in *Plasmodium* invasion, using a structure-guided strategy based on chimeric constructs combining mouse and human SR-B1 domains.

## RESULTS

### CRISPR-Cas 9 mediated inactivation of CD81 abrogates *P. berghei* infection in Hepa1-6 cells

The murine hepatoma Hepa1-6 cells express CD81 but not SR-B1 (4). In these cells, *P. berghei* sporozoite infection thus occurs via a CD81-dependent route exclusively, and can be blocked by CD81-specific antibodies or siRNA (10). To corroborate these results, we generated a Hepa1-6 cell line deficient for murine CD81 (CD81 knockout (KO) Hepa1-6 or CD81KOH16) using the CRISPR-Cas9 system. Abrogation of cell surface and total CD81 expression in CD81KOH16 cells was confirmed by flow cytometry **(Fig 1A)** and western blot **(Fig 1B)**, respectively. We then analyzed the infection phenotype of the CD81KOH16 cells using *P. berghei* sporozoites. As expected, a dramatic reduction of the percentage of *P. berghei* infected cells was observed in the CD81KOH16 cell line **(Fig 1C)**. PV quantification by microscopy after staining of UIS4, a PV membrane marker, revealed a complete inhibition of productive infection in CD81KOH16 cells **(Fig 1D)**. Intranuclear UIS4-negative parasites were observed in the CD81-deficient cells, contrasting with the well-developed EEFs with a strong UIS4 staining found in the parental Hepa1-6 cells **(Fig 1E)**. We have shown before that intranuclear parasites result from sporozoites arrested during cell traversal (11). The residual intracellular parasite population observed by flow cytometry in the KO cells **(Fig 1C)** thus likely corresponds to non-productive invasion associated with cell traversal. The CD81KOH16 cell line, which lacks CD81 and has lost susceptibility to *P. berghei* infection, thus provides a suitable tool to investigate SR-B1 function through genetic complementation experiments.

**Figure 1.**
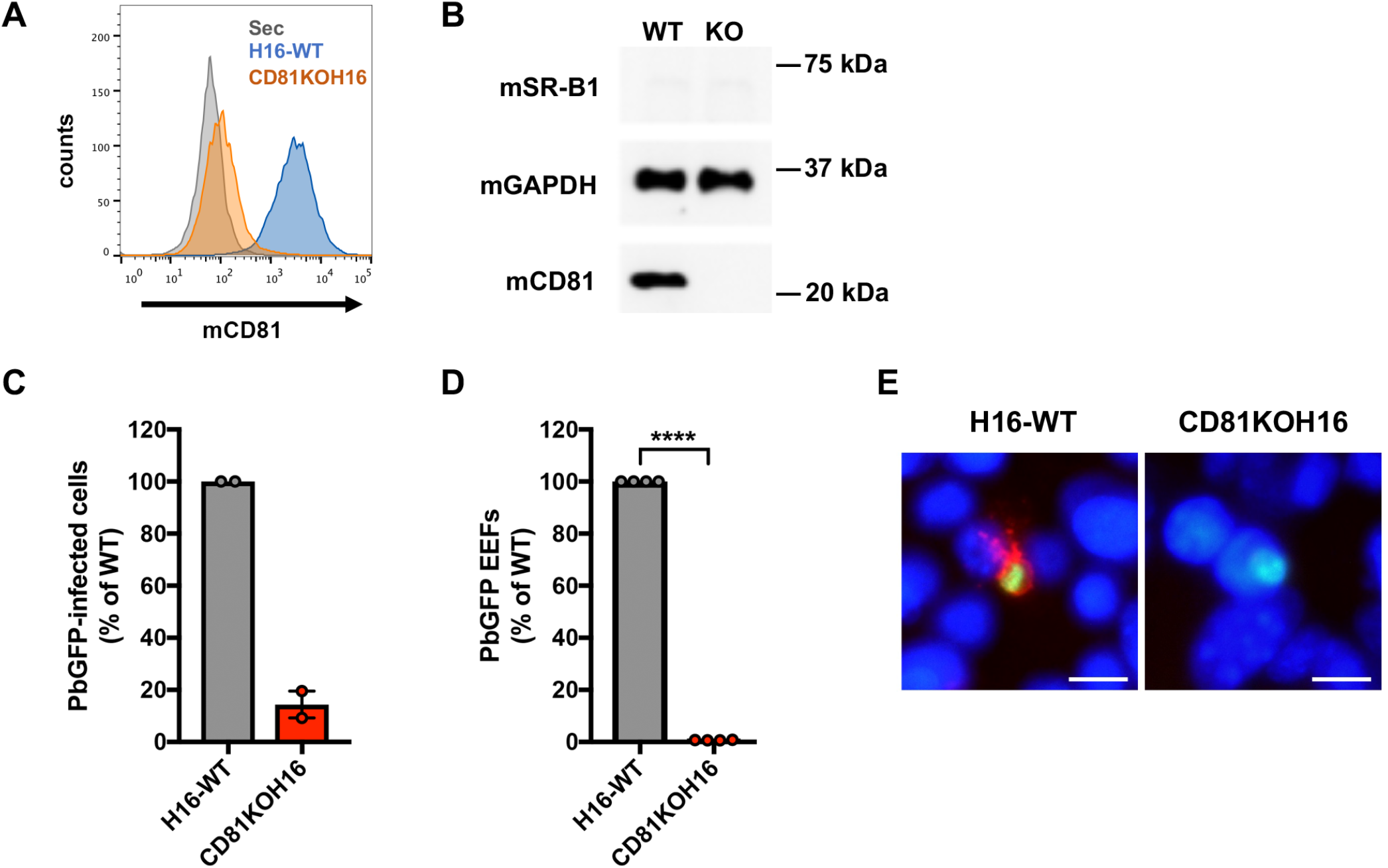
CRISPR-mediated inactivation of CD81 abrogates *P. berghei* infection in Hepa1-6 cells. **(A)** Hepa1-6 and CD81KOH16 cells were stained for surface CD81 with anti-CD81 MT81 monoclonal antibody and fluorescent secondary antibodies, before flow cytometry analysis. Histograms represent the fluorescence intensity of extracellular CD81 proteins for WT Hepa1-6 (blue) and CD81KOH16 cells (orange). The grey histogram represents cells stained with secondary antibodies only (Control). **(B)** Western blot analysis of total CD81 protein expression in WT Hepa1-6 and CD81KOH16 cells. GAPDH was used as loading control. **(C-E)** WT Hepa1-6 and CD81KOH16 cells were infected with PbGFP sporozoites and analyzed 24 hours after invasion by flow cytometry **(C)** or microscopy **(D, E)** after staining with anti-UIS4 antibodies (red) and Hoechst 33342 nuclear stain (blue). The number of EEFs per well ranged from 105 to 362 (median 175) in control cells. ****, *p*<0.0001 (ratio paired *t* test). The images show PbGFP EEFs (green) surrounded by a UIS4-positive PV membrane (red) or intranuclear parasites in CD81KOH16 cells. Scale bar, 10 μm.

### Human and murine SR-B1 differ in their ability to support *P. berghei* sporozoite infection

We have previously shown that the ectopic expression of human SR-B1 can restore *P. berghei* infection in Hepa1-6 cells where CD81 expression has been previously silenced with siRNA (4). Here, we compared the functionality of SR-B1 proteins from human and mouse origins (hereinafter referred as hSR-B1 and mSR-B1, respectively) during *P. berghei* infection after genetic complementation of CD81KOH16 cells. After transient cell transfection with plasmids encoding hSR-B1 or mSR-B1, we observed a similar expression of the two proteins by western blot **(Fig 2A)** and flow cytometry **(Fig 2B).** The transfected cells were then infected with GFP-expressing *P. berghei* sporozoites (PbGFP). In agreement with our previous observations in CD81-silenced cells (4), the transfection of hSR-B1 in CD81KOH16 cells restored their susceptibility to *P. berghei* infection **(Fig 2C)**. Unexpectedly, despite similar protein expression, mSR-B1 was not as efficient as hSR-B1 in restoring *P. berghei* infection **(Fig 2C)**. We performed similar transfection experiments in the parental Hepa1-6 cell line after CD81 silencing with siRNA, which confirmed the lower functionality of mSR-B1 protein during *P. berghei* sporozoite infection as compared to hSR-B1 **(Fig 2D)**.

**Figure 2.**
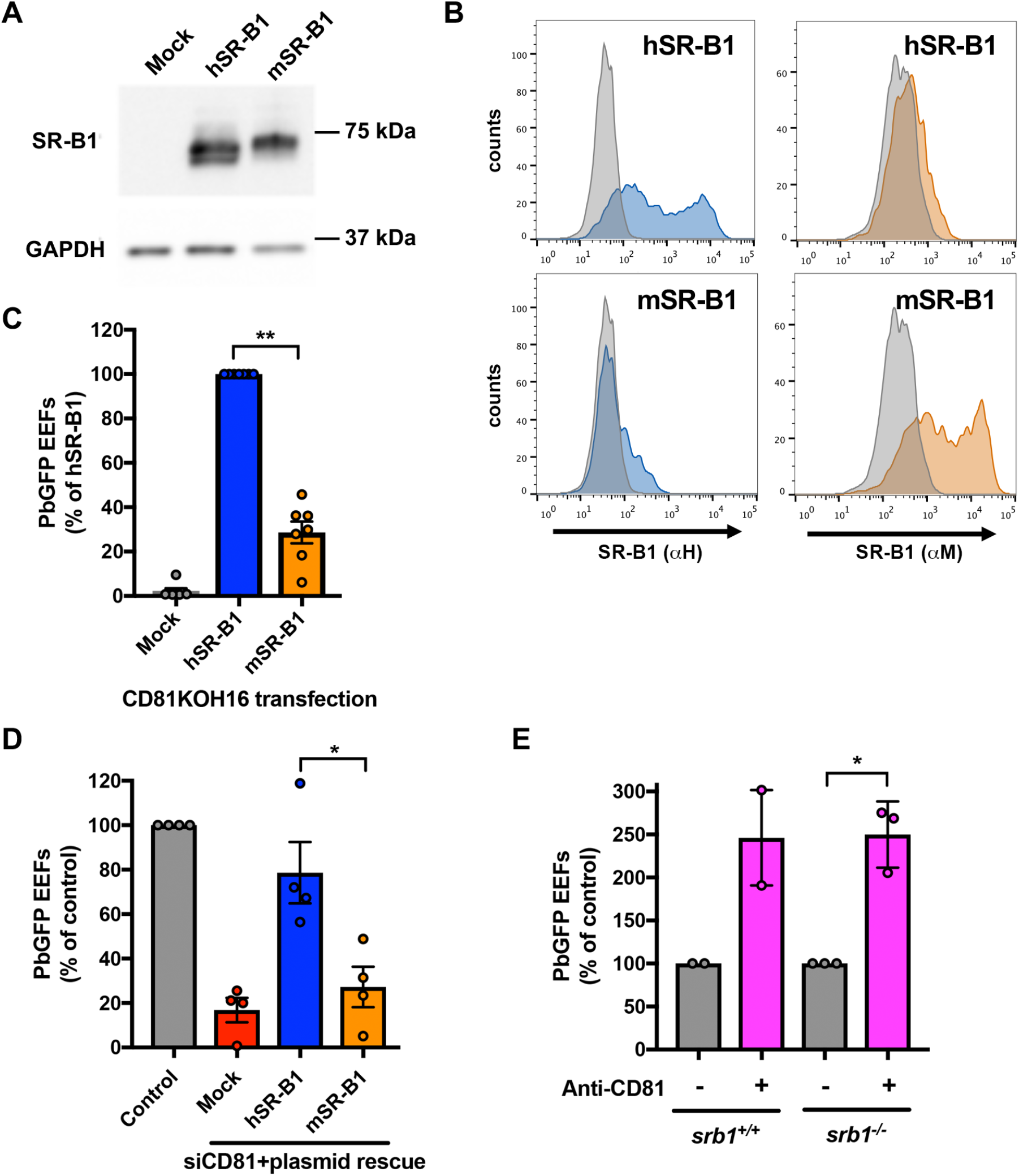
Mouse SR-B1 is poorly functional during *P. berghei* sporozoite invasion. **(A-B)** CD81KOH16 cells were transfected with either mouse or human SR-B1 plasmids, or no plasmid as a control (Mock). Total protein expression was analyzed using polyclonal anti-SR-B1 antibodies (Ab24603) by western blot **(A)** with GAPDH as a loading control. Surface protein expression was analyzed by flow cytometry **(B)** using anti-human“α-H” SR-B1 polyclonal rabbit serum (blue) and anti-mouse “α-M” polyclonal antibodies NB400-113 (orange). The grey histogram represents stained untransfected cells with the corresponding antibody. **(C-D)** CD81KOH16 **(C)** and WT Hepa1-6 cells treated with siRNA against CD81 24 hours before **(D)**, were transfected with mouse or human SR-B1 plasmids, or no plasmid as a negative control (Mock), and then infected with PbGFP sporozoites. EEFs numbers were counted by microscopy after UIS4 staining at 24 hours after sporozoite addition. The number of EEFs per well ranged from 43 to 334 (median 169) in hSR-B1-transfected cells (**C**), and from 19 to 300 (median 94) in control WT cells (**D**). *, *p*<0.05; **, *p*<0.01 (repeated measures one-way ANOVA followed by Tukey’s multiple comparisons test). **(E)** Primary hepatocytes isolated from WT or SR-B1 deficient C57BL/6 mice were infected with PbGFP sporozoites in the absence or presence of neutralizing anti-mCD81 mAb MT81, and cultured for 24 hours before EEFs quantification. *, *p*<0.05 (ratio paired *t* test).

To analyze whether the poor functionality of mSR-B1 was specific to hepatoma cells, we performed additional experiments in primary mouse hepatocytes. A previous study showed similar *P. berghei* infection rates in SR-B1^−/−^ and WT mice (2). However, the presence of a functional CD81 pathway would explain why *P. berghei* can infect the liver despite the absence of SR-B1. We thus performed infection experiments in primary hepatocytes isolated from WT or transgenic C57BL/6J mice harboring a Cre-mediated SR-B1 gene inactivation specifically in the liver (12), while using the neutralizing anti-CD81 monoclonal antibody MT81 to block the CD81 entry route (13). CD81 inhibition did not impede *P. berghei* infection of SR-B1-deficient hepatocytes, but, at the opposite, substantially increased the infection rate, similarly to WT hepatocytes **(Fig 2E)**. This enhancing effect of anti-CD81 antibodies on *P. berghei*-infection has been reported before in C57BL/6 mouse hepatocyte cultures, but the underlying mechanism remains unknown (10). Altogether, these results support the hypothesis that mouse SR-B1 does not play a prominent role during *P. berghei* sporozoite invasion in the mouse liver, and suggest that, in addition to CD81, other yet unidentified host proteins are implicated.

### Human and mouse SR-B1 protein sequence analysis and structure-homology modeling

We next investigated the structural basis that could explain the differential functionality between human and mouse SR-B1 during *P. berghei* sporozoite invasion. hSR-B1 (isoform 1) contains 509 amino acids (AA) and presents a large extracellular domain (404 AA) flanked by two transmembrane domains (both 23 AA) and two cytoplasmic tails (N-terminal: 12 AA; C-terminal: 47 AA) (6). The modeling of hSR-B1 using CD36 as a template (PDB ID: 5lgd) (7) shows that the extracellular part of the receptor can be divided into three regions: a N-terminal region (AA 36-136) harboring a thrombospondin-binding domain in the homologous CD36 protein (14), an apical region (AA 137-214) consisting of four alpha helices (α4, 5, 6 and 7), and a large C-terminal region (AA 215-439) contributing to the hydrophobic channel **(Fig 3A, C)**. The pairwise sequence alignment of hSR-B1 and mSR-B1 showed that the N-terminal and C-terminal extracellular regions were the most similar, with 81.1% and 85.7% identity, respectively, whilst the apical domain is more divergent, with 66.2% identity **(Fig 3C and D).** The hSR-B1 protein harbors 9 N-glycosylation sites, against 11 sites for mSR-B1 (15) **(Fig 3C)**. The superposition of hSR-B1 and mSR-B1 structural models revealed differences for two loops at the very top of the apex, between the α4 and α5 helices and after the α7 helix **(Fig 3B)**. We also observed differences in the electrostatic surface potentials in this area **(Fig 3E)**. When the structure is orientated in a side view to present its hydrophobic tunnel entrance, the apex lateral surface of mSR-B1 seems to be mainly electropositive whereas electronegativity is predominant in the human model **(Fig 3E)**. Remarkably, whilst the top of the apical surface is strictly neutral to electropositive in hSR-B1, mSR-B1 displays a dense electronegative region (**Fig 3E**).

**Figure 3.**
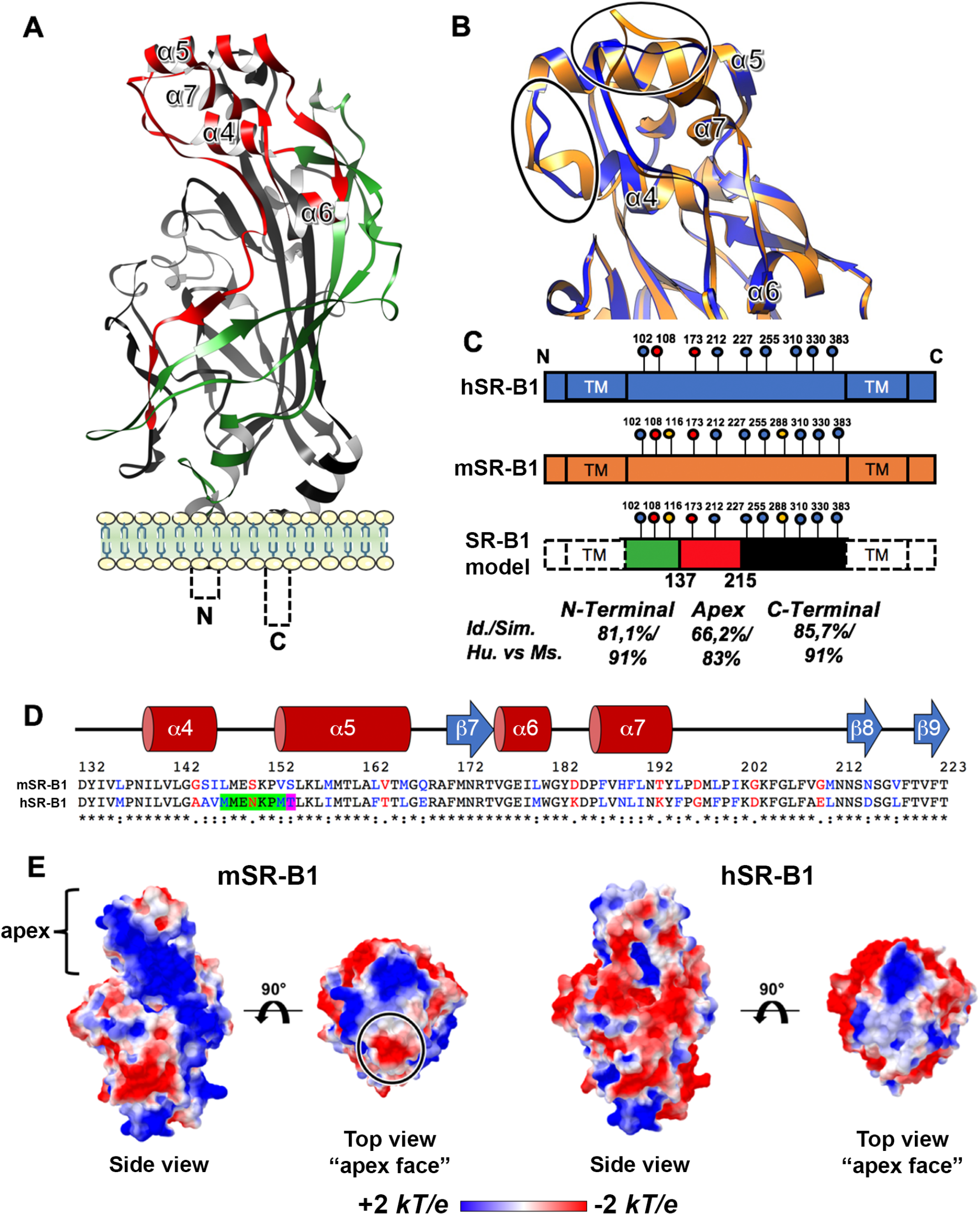
SR-B1 modeling identifies potential functional regions. **(A)** Predicted tertiary structure of hSR-B1 extracellular domain by homology modeling using CD36 (PDB ID: 5lgd) as a template, with the three regions referred to as “N-terminal” (green), “apex” (red) and “C-terminal” (black). **(B)** A close-up view of structural alignment of the apical helix bundle of mouse (orange) and human (blue) SR-B1, with their four alpha helices (α4 to α7). The main structural differences are circled in black. **(C)** Schematic representation of SR-B1 N-glycosylation sites on human (blue) and mouse (orange) proteins. Two determinant sites for SR-B1 structure and function are in red (Asn 108 and 173), mouse specific sites are in yellow (Asn 116 and 288) and conserved sites are in blue. SR-B1 model is a schematic representation of the delineated regions (“N-terminal” (green), “apex” (red), “C-terminal” (black)) in SR-B1 protein displaying all potential N-glycosylation sites. **(D)** Pairwise sequence alignment of mSR-B1 and hSR-B1 proteins for the 132-223 apical region with corresponding predicted human secondary structure (alpha helices in red and beta strand in blue). Identical, similar and different amino acids are represented in black, blue and red respectively. The threonine residue position corresponding to PfEMP1 binding phenylalanine in CD36 homolog is highlighted in purple. The residues in SR-B1 equivalent to Enterovirus-interacting site in LIMP2 are highlighted in green and purple. **(E)** Electrostatic surface potential of mSR-B1 and hSR-B1 extracellular domain from side and top views. Values are in units of kT/e at 298 K, on a scale of −2 kT/e (red) to +2 kT/e (blue). White color indicates a neutral potential. The black circle highlights a differential electrostatic surface potential between mSR-B1 and hSR-B1 at the top of the “apex” region.

### The apical domain of SR-B1 plays a crucial role during *P. berghei* infection

To determine whether the predicted structural differences at the apical domain of SR-B1 could explain the differential functionality of human and mouse SR-B1, we analyzed the functional properties of two chimeric constructs made of human and mouse sequences of SR-B1. The ApicalH chimera corresponds to a mSR-B1 backbone protein with a human apical region (AA 137-214) **(Fig 4A, B)**. Reciprocally, the ApicalM chimera corresponds to a hSR-B1 protein bearing a murine Apical region **(Fig 4A,B)**. The electrostatic surface potentials of ApicalH and ApicalM apex top are similar to human and mouse SR-B1, respectively, with only ApicalM showing a dense negatively charged region **(Fig 4C)**. CD81KOH16 cells were transiently transfected with plasmids encoding hSR-B1, mSR-B1, ApicalH or ApicalM. The two chimeras were expressed at the surface of transfected cells and detected by flow cytometry using anti-human and anti-mouse SR-B1 polyclonal antibodies **(Fig 4D)**. They were also detected by western blot analysis of whole cellular extracts **(Fig S1)**. A slightly higher band was observed in the lanes corresponding to cells transfected with mSR-B1 and ApicalH constructs a compared to hSR-B1 and ApicalM, which is likely explained by the differential glycosylation pattern of the mSR-B1 backbone **(Fig 4A)**. Cells transfected with ApicalH and ApicalM constructs bound Cy5-labelled HDLs **(Supplemental Fig S2)**, similarly to hSR-B1 and mSR-B1, suggesting that both chimeras are functional.

**Figure 4.**
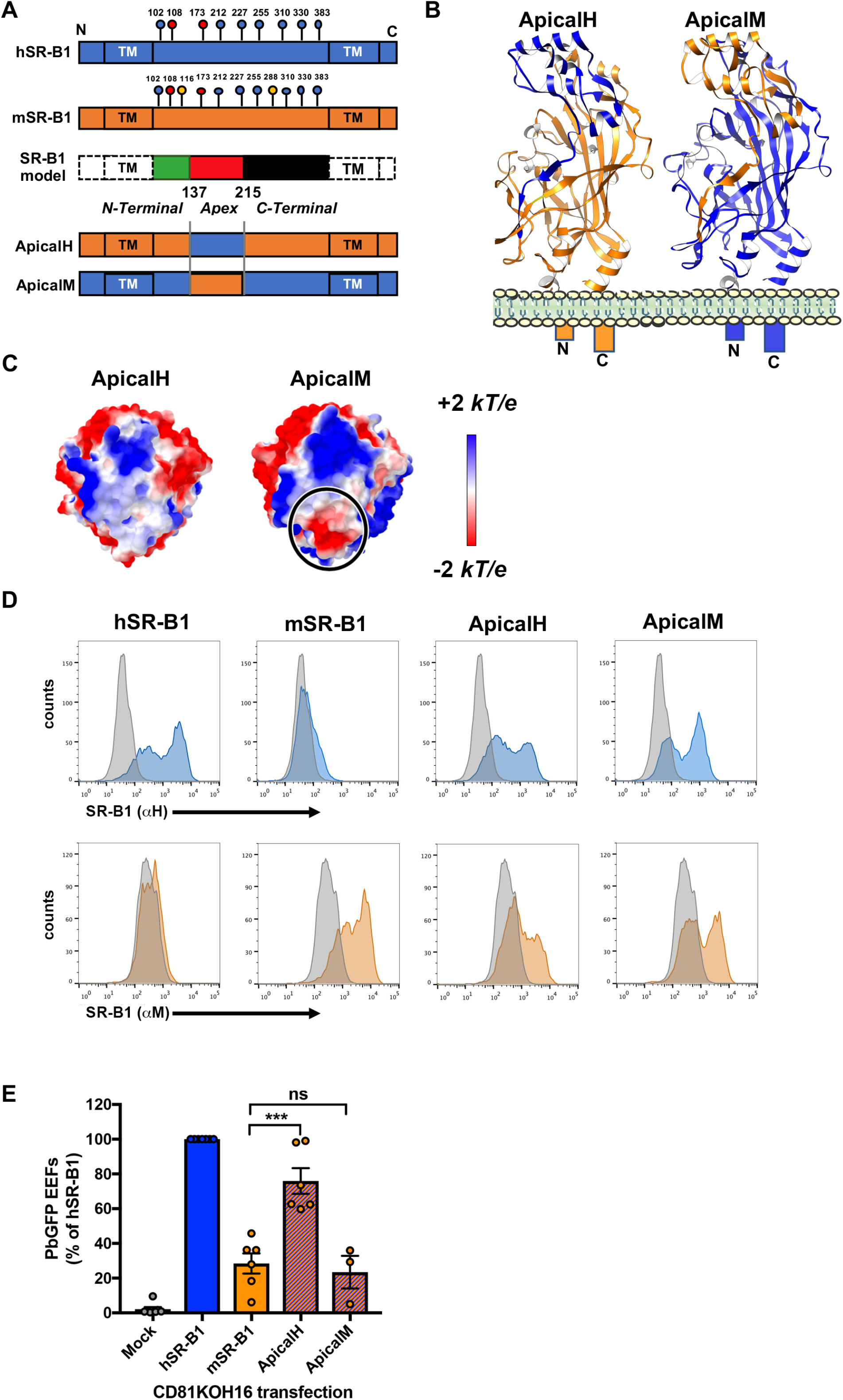
The apical domain of SR-B1 plays a crucial role during *P. berghei* infection. **(A)** Schematic representation of the ApicalH and ApicalM chimeric constructs. **(B)** Predicted tertiary structure of ApicalH and ApicalM chimeras by homology modeling, highlighting the portions of mouse (orange) or human (blue) origins. **(C)** Top views of the electrostatic surface potential of ApicalH and ApicalM chimeras’ apex. Values are in units of kT/e at 298 K, on a scale of −2 kT/e (red) to +2 kT/e (blue). White color indicates a neutral potential. The black circle highlights a differential electrostatic surface potential between the two chimeric constructs at the top of the “apex” region. **(D)** CD81KOH16 cells were transfected with hSR-B1, mSR-B1, ApicalH or ApicalM chimera plasmids, or no plasmid as a control (Mock). Protein surface expression was analyzed using anti-hSR-B1 (“αH”, blue histograms) and anti-mSR-B1 (“αM”, orange histograms), 24 hours after transfection. The grey histogram represents untransfected cells stained with the cognate antibody. **(E)** CD81KOH16 cells were transfected with hSR-B1, mSR-B1, ApicalH or ApicalM constructs, or no plasmid as a control (Mock), and infected with PbGFP sporozoites 24 hours after transfection. The number of infected cells (EEFs) was determined by microscopy after UIS4 staining at 24 hours after sporozoite addition. The number of EEFs per well ranged from 43 to 334 (median 169) in hSR-B1-transfected wells. ns, non-significant; ***, *p*<0.001 (one-way ANOVA followed by Tukey’s multiple comparisons test).

Transfected cells were then incubated with *P. berghei* sporozoites, and the number of infected cells was determined at 24 hours post-infection. These experiments revealed that replacement of the apex of mSR-B1 by that of hSR-B1 in ApicalH yielded a chimera with a 2-3 fold increase in *P. berghei* infection rates as compared to mSR-B1 **(Fig 4E)**. At the opposite, replacement of the apex of hSR-B1 by that of mSR-B1 in the ApicalM chimera resulted in a loss of function, with infection levels similar to those observed after transfection of mSR-B1 **(Fig 4E)**. Altogether, these results demonstrate that the hSR-B1 apical helix bundle (AA 137-214) is functionally determinant during *P. berghei* sporozoite invasion of hepatocytic cell lines.

### A short portion of the apical domain of SR-B1 supports *P. berghei* infection

We then sought to define more precisely the functional regions implicated in *P. berghei* infection within the apex domain. We designed three new chimeras made of a mSR-B1 backbone harboring short hSR-B1 sequences, based on both the amino acid differences between the mouse and the human sequences, and the putative interacting sites in other CD36 family receptors. The D1 chimera (AA 150-164) includes the loop between the α4 and α5 helices, where the Enterovirus 71 interacting site is located in the SR-B1 homolog LIMP-2, and encompasses a large part of the α5 helix including the PfEMP1-interacting site in CD36 **(Fig 5A-B).** The D2 chimera (AA 193-203) comprises the external tip of the α7 helix but also three phenylalanine residues in the downstream loop, exclusively present in the human sequence **(Fig 5A-B)**. The D3 chimera (AA 201-211) includes only one of these phenylalanine residues **(Fig 5A-B).** The predicted electrostatic surface potential of D1 and D3 apex top is similar to mSR-B1 **(Fig 5C),** whereas D2 apex is mostly electropositive, like hSR-B1, with no mark of electronegativity.

**Figure 5.**
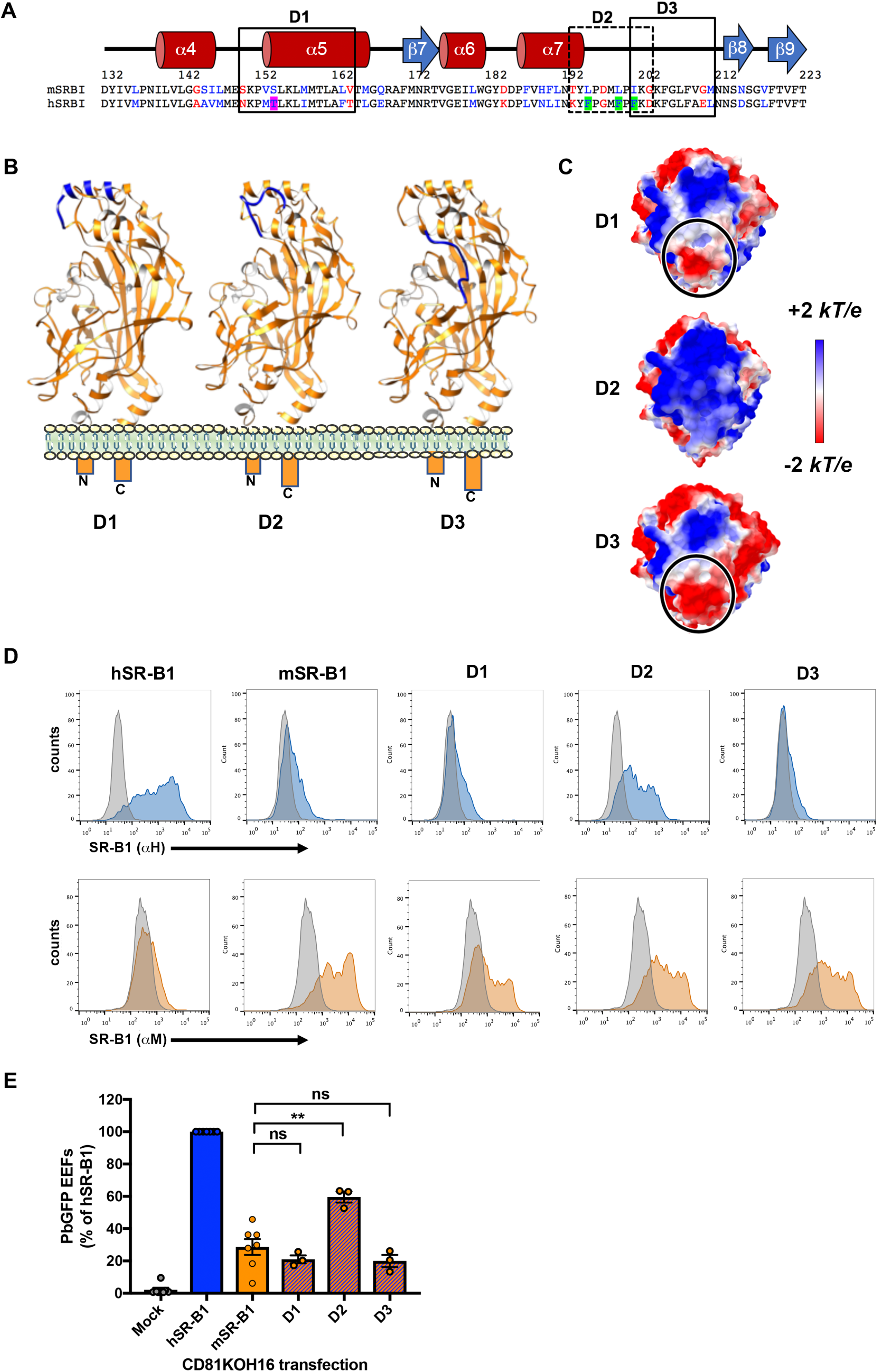
A key domain within SR-B1 apex is essential for *P. berghei* infection. **(A)** Mouse and human protein sequence alignment of the apical region AA 132-223 with the corresponding predicted human secondary structure (alpha helices in red and beta strand in blue). Identical, similar and different amino acids are represented in black, blue and red respectively. Short domains D1, D2 and D3 are delimited by boxes. **(B)** Predicted tertiary structure of D1, D2 and D3 chimeras by homology modeling, highlighting the segments of mouse (orange) or human (blue) origins. **(C)** Top views of the electrostatic surface potential of D1, D2 and D3 chimeras’ apex. Values are in units of kT/e at 298 K, on a scale of −2 kT/e (red) to +2 kT/e (blue). White color indicates a neutral potential. Blacks circle highlight a differential electrostatic surface potential between the different chimeric constructs at the top of the “apex” region. **(D)** CD81KOH16 cells were transfected with hSR-B1, mSR-B1, D1, D2, or D3 chimeric constructs. Protein surface expression was analyzed using anti-hSR-B1 (“αH”, blue histograms) and anti-mSR-B1 (“αM”, orange histograms), 24 hours after transfection. The grey histogram represents untransfected cells stained with the cognate antibody. **(E)** CD81KOH16 cells were transfected with hSR-B1, mSR-B1, D1, D2, or D3 chimeric constructs, or no plasmid as a control (Mock), and then infected with PbGFP sporozoites 24 hours after transfection. The number of infected cells (EEFs) was determined by microscopy after UIS4 staining at 24 hours after sporozoite addition. The number of EEFs per well ranged from 43 to 334 (median 169) in hSR-B1-transfected wells. ns, non-significant; **, *p*<0.01 (one-way ANOVA followed by Tukey’s multiple comparisons test).

After the transient transfection of CD81KOH16 cells, D1, D2, and D3 chimeras were all detected by flow cytometry on the cell surface using the “αM” antibody. Interestingly, only D2 was detected by the “αH” antibody, similarly to hSR-B1 and ApicalH proteins **(Fig 5D)**. Infection of the transfected cells with *P. berghei* sporozoites revealed that replacement of the AA 193-203 sequence of mSR-B1 by that of hSR-B1 in the D2 chimera resulted in a 2-fold increase in *P. berghei* infection in CD81KOH16 cells **(Fig 5E)**. In contrast, replacement of the AA 150-164 or AA 201-211 sequences in the D1 and D3 chimera, respectively, did not increase infection as compared to mSR-B1 **(Fig 5E)**. These results thus highlight the functional importance of a short 11 amino acid sequence within the hSR-B1 apical domain, which is sufficient to promote efficient *P. berghei* infection.

### The lipid transfer activity of human SR-B1 is not required for *P. berghei* infection

SR-B1 mediates selective efflux and uptake of cholesteryl esters between the plasma membrane and HDLs (16), which is mediated by SR-B1 hydrophobic channel that spans the entire length of the molecule (6). Rodrigues *et al*. reported that block lipid transport (BLT) inhibitors, which block the lipid transfer activity of SR-B1, inhibit both the entry and the development of *P. berghei* inside Huh7 cells (2). We therefore sought to determine whether in addition to the apical domain, the SR-B1 lipid transfer activity is also involved during SR-B1-dependent *P. berghei* infection of HepG2 cells. We observed an inhibition of *P. berghei* invasion in HepG2 cells when sporozoites were co-incubated with a 20 μM concentration of BLT-1 **(Fig 6A).** However, the same inhibition was also observed in Hepa1-6 cells lacking SR-B1 receptor **(Fig 6C)**. Pre-incubation of cells with BLT inhibitors before sporozoite inoculation caused no inhibition of infection in any of the cell lines tested **(Fig 6B** and **6D)**. BLT-1 at high concentration also blocked sporozoite cell traversal activity, monitored by dextran-rhodamine cellular uptake (**Fig S3**). These data strongly suggest that the inhibition caused by BLT1 is due to the toxicity of the compound on sporozoites, rather than blockage of SR-B1 function. In favor of this hypothesis, another BLT inhibitor, BLT-4, had no effect on either cell traversal or invasion by *P. berghei* sporozoites (**Fig 6A-D** and **Fig S2**).

**Figure 6.**
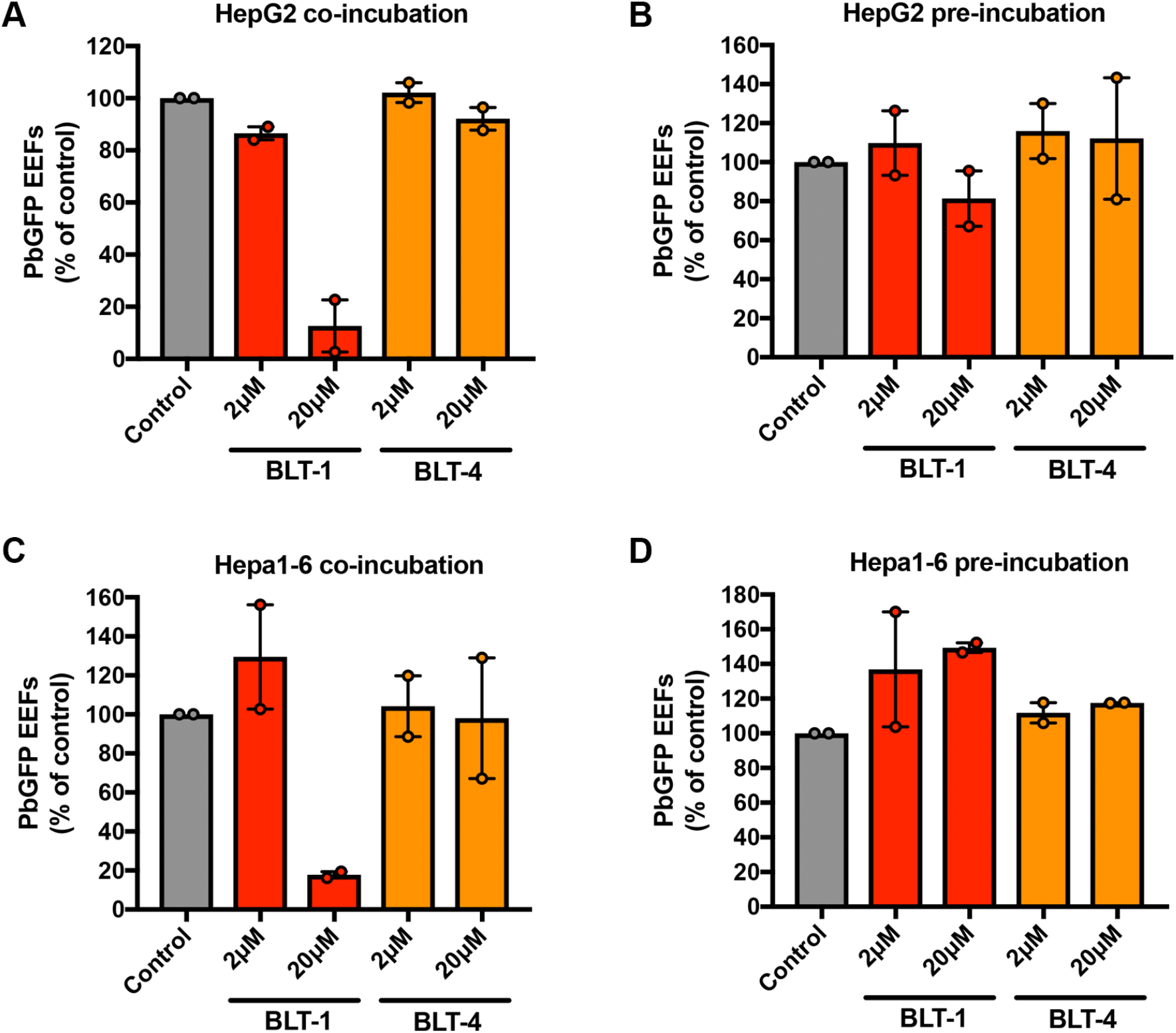
The lipid transfer activity of SR-B1 is not required during *P. berghei* infection. **(A-D)** HepG2 or and Hepa1-6 cells were treated with BLT inhibitors (BLT1 and BLT4) at two different concentrations (2 μM and 20 μM), either at the same time as sporozoite incubation (Coincubation: **A**, **C**) or prior to sporozoite addition (Preincubation: **B**, **D**). Control cells were treated with the solvant (DMSO) alone. The number of infected cells was analyzed after 24 hours by microscopy after UIS4 staining. The number of EEFs per well ranged from 223 to 545 (median 428) in control HepG2 cells, and from 43 to 378 (median 161) in control Hepa1-6 cells. All data come from two independent experiments and are represented as mean +/− range.

Altogether, these data indicate that the apical domain but not the lipid transfer activity of SR-B1 is important during *P. berghei* sporozoite entry.

## DISCUSSION

Previous studies highlighted the dual role of SR-B1 during *Plasmodium* sporozoite invasion and intracellular liver stage development (2, 5). More recently, we have shown that SR-B1 is an important host factor for *P. vivax* but not for *P. falciparum* infection, and that *P. berghei* sporozoites can use hSR-B1 as an alternative entry route to the CD81-dependent pathway (4). *P. berghei* is a rodent-infecting parasite, yet *P. berghei* sporozoites can readily infect human hepatocytic cells, using either a CD81 or a SR-B1 entry route (4). Here, we show that mSR-B1, in contrast to its human counterpart, does not support efficient *P. berghei* sporozoite invasion of hepatocytic cells. *P. berghei* was originally isolated from the African tree rat *Grammomys surdaster,* and artificially introduced for scientific purposes in the domestic mouse *Mus musculus* (17). We cannot exclude that SR-B1 function during *Plasmodium* infection may vary depending on the rodent host. In previous studies, we reported that *P. berghei* sporozoites readily infect CD81-deficient mouse hepatocytes *in vivo* and *in vitro* (3, 10), supporting the existence of alternative entry pathways. Whilst SR-B1 provides a CD81-independent route for *P. berghei* in human hepatocytic cells (4), we report here that concomitant blockage of murine CD81 and SR-B1 receptors does not prevent *P. berghei* infection in primary mouse hepatocyte cultures. These results support the existence of alternative entry routes for the parasite, which still remain to be identified. Possible candidate host receptors include the SR-B1-related proteins CD36 and LIMP-2. Although LIMP-2 is predominantly expressed in lysosomes, a fraction of the protein pool localizes at the cell plasma membrane, where LIMP-2 acts as a receptor that mediates the Enterovirus 71 host cell entry (18, 19). LIMP-2 role during *Plasmodium* infection has not been investigated so far. At the opposite, CD36 is known to play major roles during malaria infection. CD36 binds PfEMP1 variants expressed at the surface of *P. falciparum*-infected erythrocytes, and contributes to the cytoadherence of *P. falciparum* to vascular endothelial cells (20–22). It is also a major receptor for tissue sequestration of *P. berghei*-infected erythrocytes in mice (23). A previous study investigated the contribution of CD36 during *P. yoelii* and *P. berghei* sporozoite infection, using CD36-deficient mice. The data showed that both parasites could still infect hepatocytes in the absence of CD36 (24). However, in these experiments, the presence of a functional CD81-entry pathway could have masked any important role of CD36. Hence the contribution of CD36 and LIMP-2 deserves further investigation.

We took advantage of the differential functionality between human and murine SR-B1 to investigate the SR-B1 molecular determinants involved during *P. berghei* infection. Using a series of complementary chimeras designed through a structure-guided strategy, we demonstrate here the critical role of an 11 amino acid domain within hSR-B1 apical helices (AA 193 to 203) during *P. berghei* sporozoite infection. This is consistent with CD36 family proteins typically binding to a variety of ligands via their helical bundle. For instance, the ß-glucocerebrosidase binds to LIMP-2 apical domain to be delivered into the lysosome (25). Binding of Enterovirus 71 depends on a 7 amino acid sequence (AA 144-151) in LIMP-2 (18, 26). Furthermore, an apical phenylalanine of CD36 (F153) binds to *Plasmodium* PfEMP1 (7). These sites can be mapped on the SR-B1 predicted structure at the intersection between the α4 and α5 helices, at the very top of the apex. This crucial phenylalanine is replaced by a threonine in hSR-B1, and no other phenylalanine residue seems close to this area in the tertiary structure **(Fig 3D).** Interestingly, the 11 amino acid hSR-B1 functional domain we have identified contains 3 phenylalanine residues and can be mapped in the same region at the top of the apex but at a distinct site in the SR-B1 model **(Fig 3B)**.

Our data indicate that the differential activity of human and murine SR-B1 is not due to the N- or C-terminal regions of SR-B1 ectodomain, which participate in the hydrophobic channel mediating the lipid transfer. Furthermore, we did not observe any specific inhibition of SR-B1-dependent infection by BLT inhibitors. Taken together, our data suggest that the lipid transfer activity of SR-B1 is not involved during *P. berghei* sporozoite infection. Rather, we speculate that the apical helical domain of the protein may serve as a receptor for a hitherto unidentified sporozoite ligand. The dense electronegative spot at the apex of mSR-B1 and of poorly functional chimeras (ApicalM, D1 and D3), which is absent in hSR-B1 and functional chimeras (ApicalH and D2), may be unfavourable for the binding to this putative ligand. One candidate is the 6-cysteine domain protein P36, which is required for sporozoite productive invasion of hepatocytes, and is functionally linked to host receptor usage. In particular, we have shown that *P. yoelii* sporozoites genetically complemented with P36 protein from *P. berghei* can infect host cells through a SR-B1-dependent pathway (4). Whether P36 protein from *P. berghei* or from the medically-relevant *P. vivax* binds to the apical helix bundle of SR-B1 remains to be determined.

In conclusion, this study provides new insights into the function of SR-B1 during malaria infection, and paves the way towards a better characterization of the molecular interactions leading to parasite entry into hepatocytes. Our results may be particularly relevant to *P. vivax* malaria, as SR-B1 is the first and up to now only known host entry factor for *P. vivax* sporozoites (4). The characterization of SR-B1 molecular function and the identification of interacting parasite ligands may lead to the development of novel intervention strategies to prevent *P. vivax* sporozoite entry, before the establishment of the liver stage and the hypnozoite reservoir.

## MATERIALS AND METHODS

### Ethics statement

All animal work was conducted in strict accordance with the Directive 2010/63/EU of the European Parliament and Council ‘On the protection of animals used for scientific purposes’. Protocols were approved by the Ethical Committee Charles Darwin N°005 (approval #7475-2016110315516522).

### Experimental animals, parasite and cell lines

We used GFP-expressing *P. berghei* (PbGFP, ANKA strain) parasites, obtained after integration of a GFP expression cassette at the dispensable p230p locus (27). PbGFP blood stage parasites were propagated in female Swiss mice (6–8 weeks old, from Janvier Labs). *Anopheles stephensi* mosquitoes were fed on PbGFP-infected mice using standard methods (28), and kept at 21°C. PbGFP sporozoites were collected from the salivary glands of infected mosquitoes 21–28 days post-feeding. Hepa1-6 cells (ATCC CRL-1830) and HepG2 (ATCC HB-8065) were cultured at 37°C under 5% CO2 in DMEM supplemented with 10% fetal calf serum (10500064, Life Technologies), L-glutamine 20 μM (25030024, Life Technologies), and 1% penicillin-streptomycin, as described (10). Primary mouse hepatocytes were isolated by collagenase perfusion (C5138, Sigma), as described in (29), from C57BL/6 mice harboring a Cre-mediated SR-B1 gene inactivation specifically in the liver (12). Primary hepatocytes were seeded at confluency in 96 well plates and cultured at 37 °C in 4% CO2 in William’s E medium (22551022, Life Technologies) with 10% fetal calf serum, 1% penicillin-streptomycin (15140122, Life Technologies), hydrocortisone 50 μM (Upjohn laboratories SERB) and 1% L-glutamine, Bovine insulin 5 μg/ml (I5500, Sigma) for 24 hours before sporozoite infection.

### Small interfering RNA silencing of CD81

The siRNA oligonucleotide against CD81 (5’-CGUGUCACCUUCAACUGUA-3’) was validated in previous studies (10). Transfection of siRNA oligonucleotides was performed by electroporation in presence of 10 μL of siRNA 20 μM, as described (11). Cells were cultured during 48 hours before infection or analysis by immunofluorescence. As negative controls, we used cells electroporated in the absence of siRNA oligonucleotide.

### Generation of a CD81KOH16 cell line using CRISPR-Cas9

The day before transfection, Hepa1-6 cells were plated in 24 well plates at a density of 90 000 cells per well. Cells were transfected with 500 ng of LentiCrispR V2 (Addgene plasmid #52961) containing a guide RNA targeting mouse CD81 (GCAACCACAGAGCTACACCT) using Lipofectamine 2000 (11668027, Life Technologies). Puromycin selection was carried out 36 hours after transfection using a 5 μg/ml solution. Cells were exposed to puromycine for 48 hours, then washed and expanded for two weeks in complete medium before analysis. Immunostaining was performed using the rat monoclonal antibody MT81 to label mouse CD81 (Silvie et al., 2006a). All incubations were performed at 4°C during one hour. We used AlexaFluor-488 Goat anti-rat antibody (A1106, Life technologies) as a secondary antibody. Cells were then fixed with 1% formaldehyde solution and analyzed using a Guava EasyCyte 6/2L bench cytometer equipped with 488 nm and 532 nm lasers (Millipore).

### Homology modeling of SR-B1 chimeras

The SR-B1 amino acid sequence of *H. sapiens* (Uniprot: Q8WTV0) was submitted to the HHpred interactive server for remote protein homology detection (30). The server identified the X-ray structure of the scavenger receptor CD36 (PDB ID: 5lgd) at 2.07 Å resolution (7) as the best template to model the SR-B1 protein (probability: 100%, e-value: 2.3e-91). Sequences of SR-B1 chimeras were aligned and modeled using Swiss-Model through the ExPAsy molecular biology suite (31). Each SR-B1 model was then subjected to loop refinement and energy minimization using GalaxyRefine (32) and YASARA (33), respectively. SR-B1 models were validated for quality using MolProbity for local stereochemistry (34), and Prosa II for global 3D quality metrics (35). Additionally, we validated the structure by checking that all the N-glycosylation sites were solvent-exposed.

The protein electrostatic surface potential was calculated using Adaptive Poisson-Boltzmann Solver (APBS) (36), after determining the per-atom charge and radius of the structure with PDB2PQR v.2.1.1 (37). The Poisson-Boltzmann equation was solved at 298 K using a grid-based method, with solute and solvent dielectric constants fixed at 2 and 78.5, respectively. We used a scale of −2 kT/e to +2 kT/e to map the electrostatic surface potential in a radius of 1.4 Å. All molecular drawings were produced using UCSF Chimera (38).

### SR-B1 chimeric construct design and plasmid transfection

Plasmids encoding human and mouse SR-B1 have been described previously (39, 40). The ApicalH and ApicalM chimera were obtained by cloning a single insert amplified from chimeric synthetic genes (Eurofins Genomics) into the mSR-B1 and hSR-B1 plasmids, respectively. The D1, D2 and D3 chimera were generated by inserting into the mSR-B1 plasmid two fragments amplified with primers containing hSR-B1 sequences. The sequence of all oligonucleotides used to amplify DNA inserts and the sequence of synthetic genes used as templates are indicated in Supplemental Table 1. Information on plasmid sequence is available on demand. All cloning steps were performed using In-fusion cloning kit (639649, Ozyme) and controlled by Sanger sequencing (Eurofins genomics). High concentration plasmid solutions were produced using XL1-Blue Competent Cells (200249, Agilent technology) and plasmid extraction was performed using Qiagen Plasmid Maxikit (12163, QIAGEN) according to the manufacturer’s recommendations. Transfection of SR-B1 or chimeras encoding plasmids was performed 24 hours after siRNA electroporation, or directly on CD81KOH16 cells, using the Lipofectamine 2000 reagent (11668027, Life Technologies) according to the manufacturer’s specifications. Following plasmid transfection, cells were cultured for an additional 24 hours before sporozoite infection or protein expression analysis.

### Western blot

After cell lysis in 1% NP-40, soluble fractions were analyzed by western blot under non-reducing conditions, using a Biorad Mini-Protean® electrophoresis chamber for SDS-PAGE and transfer on polyvinylidene fluoride (PVDF) membranes. Membranes were probed with anti-mouse CD81 MT81 (13) at 2 μg/ml, anti-mSR-B1 polyclonal antibody (Ab24603) diluted at 0.9 μg/ml, and anti-mouse GADPH (TAB1001) as a loading control (0.5 μg/ml). Chemiluminescence detection was performed using ECL Prime reagents (RPN2232,GE healthcare Life sciences) and an ImageQuant LAS 4000 system (GE Healthcare).

### Immunofluorescence assays

For the immunolabeling of SR-B1 and chimeric proteins, cells were harvested using an enzyme-free Cell Dissociation buffer (13151014, Thermofisher). All incubations were performed at 4°C in PBS/BSA 3% during one hour with either “αH” anti-SR-B1 polyclonal rabbit serum (40) or “αM” anti-SR-B1 polyclonal rabbit antibodies NB400-113 (Novus Biological). We used AlexaFluor-488 Donkey anti-rabbit antibody (Ab150073, Life technologies) as secondary antibody with a 45 minutes incubation. After fixation in 1% formaldehyde, cells were analyzed using a Guava EasyCyte 6/2L bench cytometer equipped with 488 nm and 532 nm lasers (Millipore). Flow cytometry plots are representative of at least three independent experiments.

### In vitro infection assays

Hepa1-6 cells were seeded in 96 well plates (2×10^4^ per well seeded the day before transfection) and incubated with 1×10^4^ PbGFP sporozoites for 3 hours, washed, and further cultured until 24 hours post-infection. HepG2 and HepG2/CD81, plated in 96 well plates with 3×10^4^ cells per well seeded the day before infection, were infected using 5×10^3^ PbGFP sporozoites. In some experiments, anti-mouse CD81 MT81 at 20 μg/ml (13), BLT-1 (SML0059, Sigma), BLT-4 (SML0512, Sigma) (both prepared in pure DMSO) or diluted DMSO, were added to sporozoites during infection. For the dextran assay, 0.5 mg/ml rhodamine-conjugated dextran (Life technologies) was added to sporozoites during infection. Infected cultures were then either trypsinized for detection of GFP-positive cells and/or dextran-positive cells by flow cytometry on a Guava EasyCyte 6/2L bench cytometer (Millipore), or fixed with 4% paraformaldehyde and analyzed by fluorescence microscopy after labeling with antibodies specific for UIS4 (Sicgen) and the nuclear stain Hoechst 33342.

### HDL binding assay

Human HDL lipoproteins (LP3, Calbiochem) were labeled using the Cy5 monoreactive Dye pack (PA25001, GE Healthcare) and filtered using Illustra microspin G25 columns (27532501, GE Healthcare). CD81KOH16 cells were dissociated at 24 hours post-transfection using an enzyme-free Cell dissociation buffer (13151014, Thermofisher) and incubated with Cy5 labeled HDLs (5 μg/ml) for 20 minutes at 37°C. After washing, they were incubated with Suramin (574625, Merck Millipore) at 10mg/ml for one hour at 4°C. HDL binding to SR-B1 and chimeras was then evaluated by flow cytometry using the BD LSR Fortessa™.

### Statistical analyses

Statistical analyses were performed with GraphPad Prism on at least three independent experiments, each performed in triplicates, as indicated in the legend to the figures. All graphs show the mean ± SEM (unless otherwise indicated) expressed as percentage of control (WT cells or CD81KOH16 cells transfected with hSR-B1, as indicated).

## ACKNOWLEDGMENTS

We thank Jean-François Franetich, Maurel Tefit, Mariem Choura and Thierry Houpert for rearing of mosquitoes and technical assistance, and Drs Maryse Lebrun and Jérôme Clain for fruitful discussions. This work was funded by the European Union (FP7 PathCo Collaborative Project HEALTH-F3-2012-305578), the Laboratoire d’Excellence ParaFrap (ANR-11-LABX-0024), and the Agence Nationale de la Recherche (ANR-16-CE15-0004). GM was supported by a “DIM Malinf” doctoral fellowship awarded by the Conseil Régional d’Ile-de-France.

## FIGURE LEGENDS

**Supplemental figure 1.**
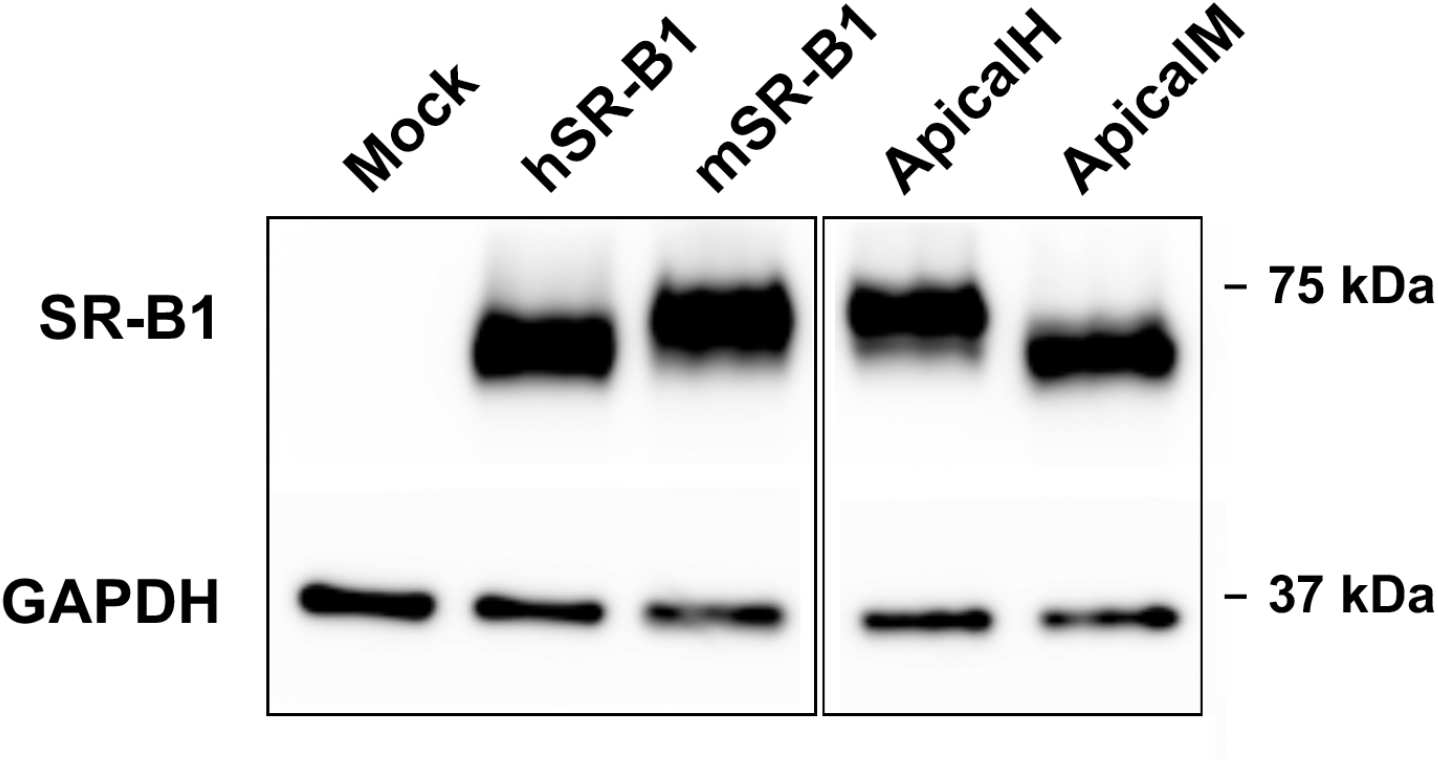
CD81 KO Hepa1-6 cells were transfected with either mSR-B1, hSR-B1, ApicalH or ApicalM construct plasmids, or no plasmid as a control (Mock). Total protein expression was analyzed by western blot using polyclonal anti-SR-B1 antibodies (Ab24603) and anti-GAPDH antibodies as a loading control.

**Supplemental figure 2.**
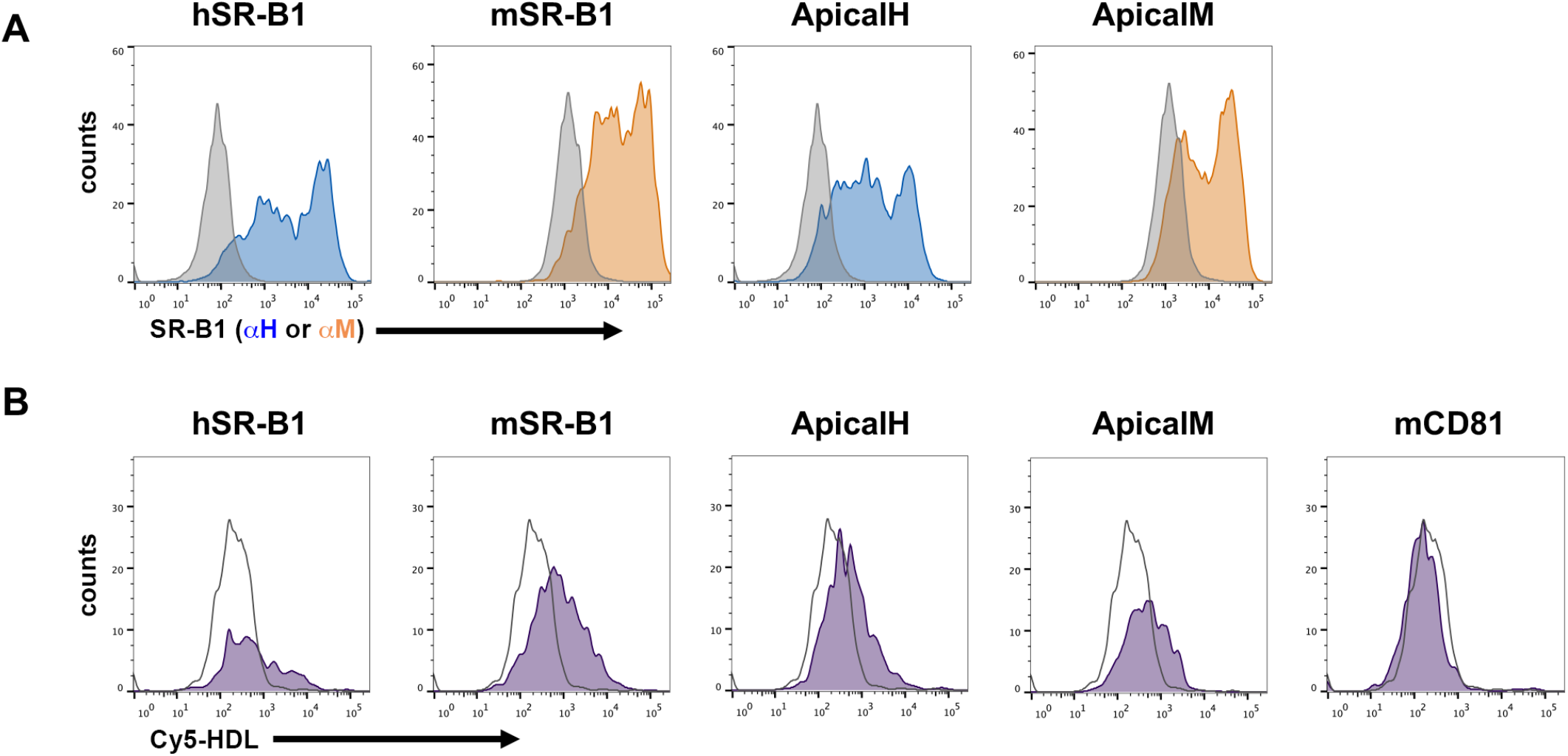
**(A-B)** CD81KOH16 cells were transfected with hSR-B1, mSR-B1, ApicalH or ApicalM chimeric constructs, or with a plasmid encoding mCD81 (negative control). **(A)** Protein surface expression was analyzed using anti-hSR-B1 (“αH”, blue histograms) and anti-mSR-B1 (“αM”, orange histograms), 24 hours after transfection. The grey histogram represents untransfected cells stained with the corresponding antibody. **(B)** Cy5 fluorescent HDLs were added to transfected cells 24 hours after transfection to measure HDL binding (purple peak). HDL binding to non-transfected cells (negative control) is shown as a white peak. Cells transfected with a mouse CD81 construct did not bind HDLs, as expected.

**Supplemental figure 3.**
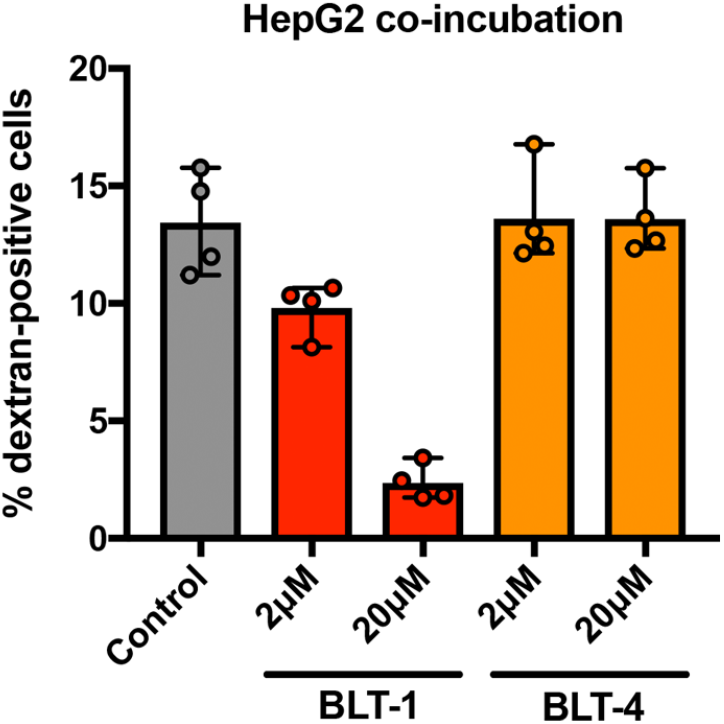
HepG2 cells were infected with PbGFP sporozoites in the presence of rhodamine-conjugated dextran and BLT inhibitors (BLT1 and BLT4) at two different concentrations (2 μM and 20 μM), or DMSO as a control. Dextran-positive cells were analyzed by flow cytometry 3 hours after sporozoite addition.

**Supplemental table 1.**
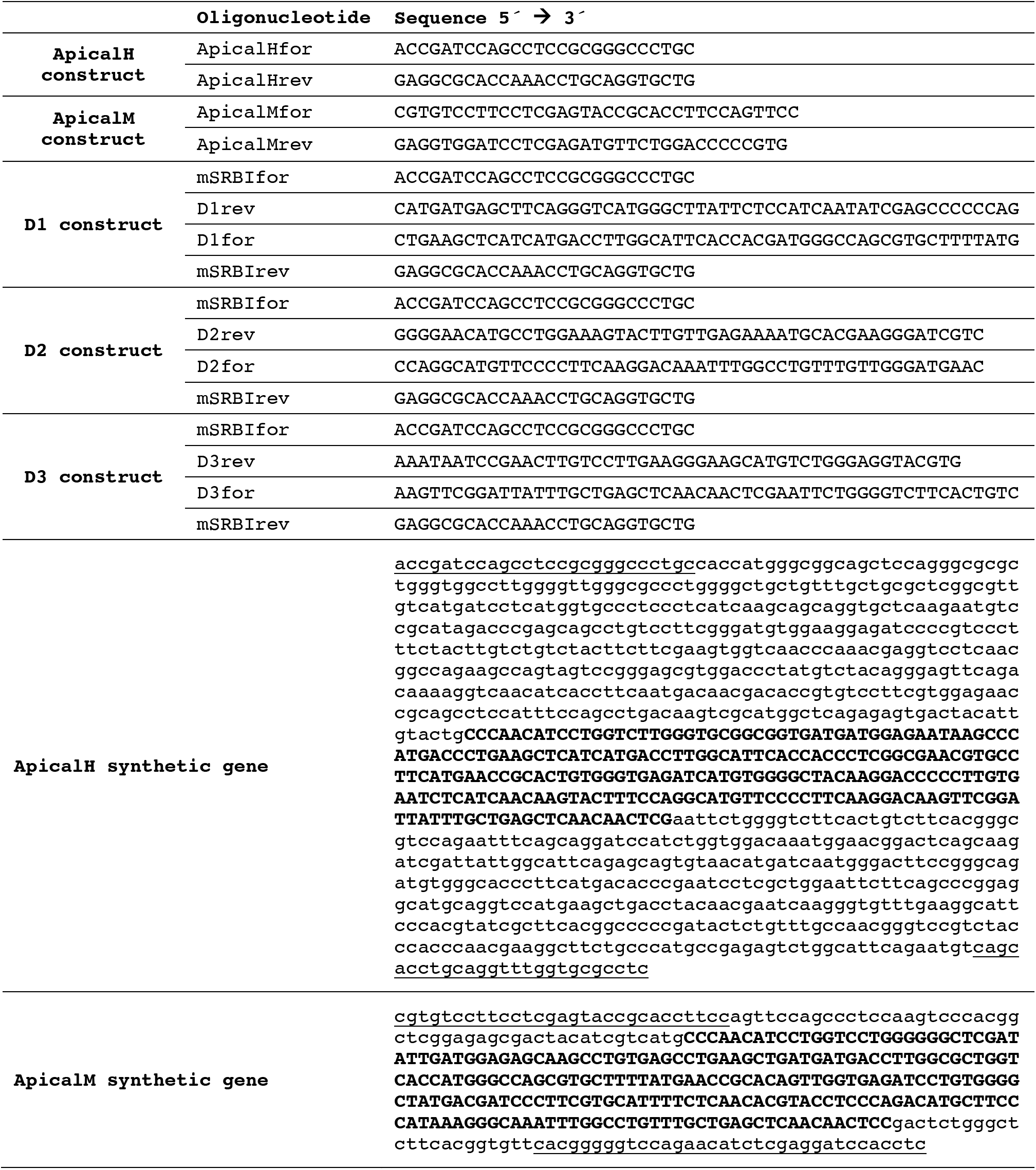
Sequences of oligonucleotides and synthetic genes used in this study.

